# A mixed model approach for estimating drivers of microbiota community composition and differential taxonomic abundance

**DOI:** 10.1101/2020.11.24.395715

**Authors:** Amy R Sweeny, Hannah E Lemon, Anan Ibrahim, Kathryn A. Watt, Kenneth Wilson, Dylan Z Childs, Daniel H Nussey, Andrew Free, Luke McNally

## Abstract

1. Next-generation sequencing (NGS) and meta-barcoding approaches have revolutionized understanding of within-host communities, such as the gut microbiome, in humans and laboratory animals. The application of such approaches in wild animal populations is growing, but there is a disconnect between the widely-applied generalised linear mixed model (GLMM) approaches commonly used to study phenotypic variation and the statistical toolkit from community ecology which is typically applied to meta-barcoding data.
2. Here, we describe and illustrate a novel GLMM-based approach for analysing the taxon-specific sequence read counts derived from standard meta-barcoding data. This approach allows us to decompose the contribution of different drivers of variation in community structure (e.g. year, season, individual host), via interaction terms in the random effects structure of the model. We also show how these models can be used to determine the degree to which specific taxa or taxonomic groups are responsible for variance attributed to different drivers.
3. To illustrate this approach, we applied it to two cross-sectional meta-barcoding data sets from the Soay Sheep population of St. Kilda. The GLMM approach yielded results that were in agreement with more classical approaches from community ecology, showing that variation the gut microbiota community in these sheep was better explained by age group than by season. We were able to quantify the contributions of different sources of variation to community structure, and also to drill down into the model predictions to show that the age effects we observed were principally due to increases in taxa of the phyla Bacteroidetes and declines in taxa of the phyla Firmicutes.
4. Our proposed models offer a powerful new approach to understanding the drivers of variation in estimates of community structure derived from meta-barcoding data. We discuss how our approach could be readily adapted to allow researchers to estimate that contribution of host genotype, environment, and microbial/parasite phylogeny to observed community structure, and thus provide a powerful means to answer emerging questions surrounding the ecological and evolutionary roles of within-host communities.

## 1. Introduction

The ecological dynamics of within-host communities of parasites and commensal microbes can have dramatic effects on host health and fitness (Alberdi *et al*. 2016; Koskella *et al*. 2017). One increasingly well studied example of such a within-host community is the so-called gut microbiota: the often complex and diverse community of commensal bacteria resident to the gastrointestinal tracts of their animal hosts. As well as playing a crucial role in the digestion of food, studies from humans and model laboratory animals highlight impacts of the gut microbiota on host behaviour, metabolism, as well as endocrine and immune homeostasis (Sudo *et al*. 2004; Cox-Foster *et al*. 2007; Desbonnet *et al*. 2014; Wahlström *et al*. 2016). A growing number of studies within ecology and evolutionary biology are seeking to study the dynamics of the gut microbiota of natural systems using a combination of faecal sampling and next-generation sequencing (NGS) meta-barcoding approaches. Understanding the role of within-host communities in underpinning host phenotypic variation, as well as wider ecological and evolutionary dynamics, in the wild will require statistical approaches that allow us to robustly quantify the contribution of different environmental and host-related factors to such metabarcoding data. Generalised linear mixed models (GLMM) are a well-established and widely used suite of statistical models within ecology and evolution which provide a flexible means for appropriately dealing with the complex data structures and relationships between predictors of interest that arise in ecological data (Bolker *et al*. 2009). Although they have yet to be widely applied in this context, GLMMs have huge potential to help dissect and understand the drivers of within-host community dynamics, as revealed by meta-barcoding data.

Standard methodologies for investigating hypotheses concerning gut microbiota dynamics in the wild typically include collection of faecal samples from selected study subjects and application of next-generation sequencing (NGS) techniques for meta-barcoding of informative bacterial genes for taxonomic assignment of sequenced reads (J *et al*. 2018). Microbiota community analysis commonly relies on transformation of OTU (operational taxonomic units) or ASV (amplicon sequence variant) counts into relative proportions per sample or rarefaction such that a set library size is randomly subsampled from all samples (McMurdie & Holmes 2014; Weiss *et al*. 2017; McMurdie 2018) Hypothesis testing using transformed counts from 16S taxonomic assignments typically is focused on community-level differences in taxonomic diversity and composition between experimental groups or time points of interest. Statistical approaches to this end include estimation of alpha diversity (the number of distinguishable taxa within a sample), distance measures (e.g. Bray-Curtis dissimilarity (Bray and Curtis, 1957)), and ordination with dimensionality reduction (e.g. principal coordinates analysis). Data transformations and hypothesis tests in these approaches have several limitations. Standardisations of data based on proportions ignore heteroscedasticity from different library sizes across samples, while those relying on rarefaction restrict data such that the reads considered per each sample are limited to the minimum number of reads across all samples (McMurdie & Holmes 2014). This in turn can significantly elevate rates of false positives or reduce performance in microbiome clustering approaches. In addition to statistical pitfalls, these traditional approaches for assessing community-level differences differ philosophically from GLMM-based approaches which partition complex sources of variance. Although traditional approaches have provided substantial insights into microbiota community composition, they fall short of the flexibility and power offered by GLMM-based approaches to dissect the manifold and complex contributors to variation in measured phenotypes in natural populations. There has therefore been movement among community ecologists toward such model-based methods (Niku *et al*. 2019). Here we develop a GLMM-based approach to decompose the sources of variation in count data derived from meta-barcoding approaches and discuss the advantages of this approach for analysis of microbiota and other community data.

The application of mixed effects models to microbiota datasets is not new. The “Hierarchical Modelling of Species Communities” (HMSC) approach uses latent variable modelling and random effects to model community compositions 1 and has been applied to the composition of the microbiota (refs). A similar approach developed for microbiota datasets has also shown insights into microbiota composition in the wild (ref Bjork). Our suggested approach differs from these in several ways. Firstly, these approaches have a focus on modelling correlations among microbial taxa using latent variables to model residual correlation, which adds quite a lot of complexity to the modelling process. Our approach does not attempt to model these correlations, it focuses on variance decomposition of the sort familiar to ecologists and evolutionary biologists working on wild systems. If correlations among microbial taxa is of primary interest we would direct readers to these approaches. Secondly, our approach does not require the use of any particular modelling package or a high degree of proficiency in coding. There are two central ideas in our approach – using sample level random effects in Poisson models to account for variability in library size, and using random effects of microbial taxonomy to allow for effects of host and environment on microbiota composition – that can be implemented in almost any random effects modelling software or packages with which the reader is familiar. Thus, at the expense of modelling residual correlation among species, our approach offers a familiar method to decompose sources of variance in the microbiota for field scientists. Below we outline the motivation for this approach and illustrate this via an application to two 16S metabarcoding datasets from a wild mammal.

## 2. A GLMM approach

As gut microbes have such important effects on host physiology, behaviour and health much research has sought to identify individual microbial taxa that are responsible for alterations of host phenotype and state. This has in many ways mirrored the goals of many genome-wide association study (GWAS) analyses, which have sought to identify particular genetic variants associated with phenotypes of interest, often with a goal of developing diagnostics or drug targets (Visscher *et al*. 2012). However, just as GWAS analyses have shown us that most phenotypes are highly polygenic, being determined by a complex combination of genetic variants of small effects (Goldstein 2009; Consortium 2009; Loh *et al*. 2015), the study of host-associated microbiomes has often failed to find single taxa associated with host states (Clemente *et al*. 2012; Vayssier-Taussat 2014). Instead many changes in host state are associated with general shifts in microbiome composition, often termed dysbiosis (Clemente *et al*. 2012; Carding *et al*. 2015). Phenomena such as dysbiosis shift the level at which we look for associations with host phenotype move from a small number of microbial taxa to the whole microbiota. In addition, the most pressing questions about host-associated microbiota in ecology and evolution are very general and focused on the entire microbiota community (Koskella *et al*. 2017). For example, what are the relative roles of host physiology and environment in shaping the microbiota? How heritable is the microbiota? How much does microbiota composition impact fitness? The shift in focus of these questions from individual taxa to complete community poses an important conceptual and statistical challenge.

As previously discussed, host-associated microbiotas often constitute hundreds or thousands of different taxa. Whenever we need to estimate a large ensemble of related parameters, a common statistical approach is to treat them as random variables from some distribution (Gelman & Hill 2006). To understand how this approach applies to the microbiota let us consider the concrete question of estimating how the composition of the gut microbiota might change with season in a wild mammal. In traditional approaches to analysing microbiota datasets, it would be common to visualise an ordination of the data, distinguishing points by season. Then, one would perform a permutational ANOVA on a dissimilarity matrix to test if microbiotas from different seasons are more dissimilar than those from the same season, and go on to test for differential abundance of individual taxa across seasons to identify taxa with a major role in these changes (Segata *et al*. 2011; Mandal *et al*. 2015). In this approach, estimates for how individual taxa differ by season are all independent of each other. Using a random effects model would approach this in a fundamentally different manner, where a model would be fitted with a random effect for the seasonal change for each microbial taxon. This approach has the advantage that all taxa inform the estimate of the mean and variance of the distribution that the effects across taxa come from. The estimates of parameters for individual taxa are then “shrunk” to this distribution. This “shrinkage” is known to improve the accuracy of parameter estimation as long as there are large numbers of groups for the random effects, which is generally true for most host-associated microbiota owing to their large number of taxa.

While fitting such random effects models is known to improve parameter estimation owing to shrinkage, its biggest advantage is in allowing us to shift the questions we ask to the whole microbiota level, and partition complex & inter-related sources of variance. GLMM approaches have been used across other ecological and evolutionary contexts to estimate repeatability, relative levels of spatiotemporal variance (Albery *et al*. 2019), social and common environment effects (Rushmore *et al*. 2013; Froy *et al*. 2018), as well as heritability and the role of host genetics (Hayward *et al*. 2014). Answering such questions has proved hugely challenging in the microbiota field as most analyses rely on tools which are not multi-level, from which it is extremely difficult to decompose the relative contribution of simultaneous processes at the host and environmental scale. However, mutli-level models have been shown to offer significant advantages over many other compositional methods in community ecology for species abundance data (Jackson *et al*. 2012). Here, we develop and illustrate on method to appropriately structure random effects across microbial taxa within a community using a GLMM, and thus partition the sources of variance driving microbiota composition.

To see how such a model can be structured let us again return to the example of estimating the effect of season on the gut microbiota of a wild mammal (Figure 1). Consider a scenario with two samples taken per host from a sample of hosts in a population, one in winter and one in summer, and with samples appropriately sequenced and reads bioinformatically assigned to ASVs. This will yield data in the form of a count of reads belonging to each ASV (the focal taxonomic group) within each sample (Figure 1A), with two samples per individual host one from each season. We can directly analyse such count data by fitting a Poisson family GLMM with log-link. The predicted values on the link scale are given according to the following model (Figure 1B).

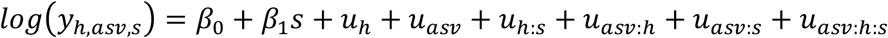

**Figure 1.**
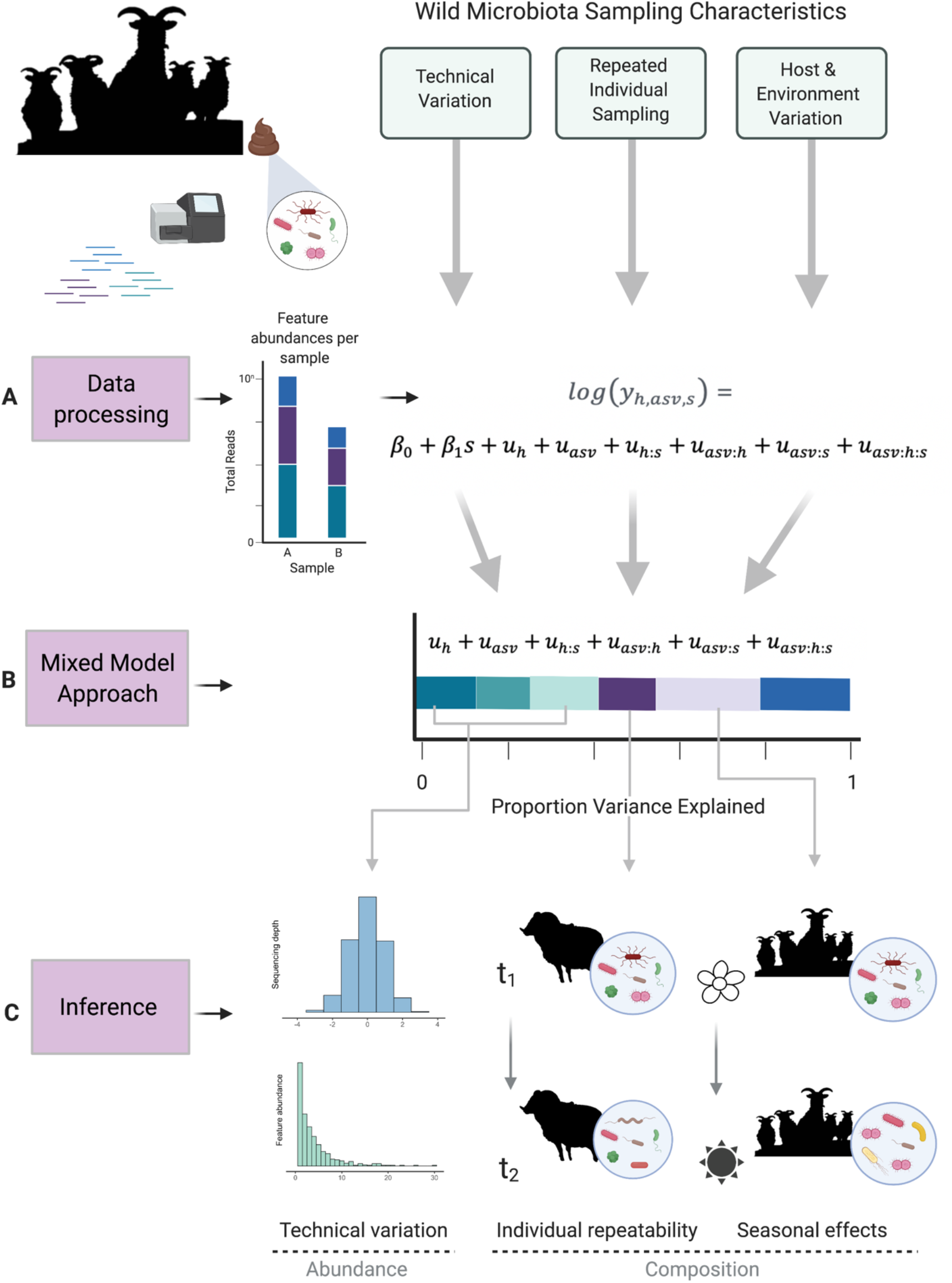
Overview of mixed model approach to wild microbiota analysis. Data processing (A) generates amplicon sequence variant (ASV)-level abundances for each sample. These raw abundances are used as the response for generalised linear mixed-effects models with Poisson error families. In the example illustrated, data includes sampling timepoints for a group of individuals taken during two seasons. Model syntax therefore specifies a fixed effect of age, and random effects for taxonomy (asv), sample id (host:season, h:s), individual differential abundance of asvs (asv:h), differential abundance of asvs across seasons (asv:s), and a residual variance at the row-level (asv:h:s). GLMM output can be used to partition the variance explained by each random effect term (B). These variance components can be interpreted as the relative contributions of both technical variation and host or environmental contributions to differential abundance as illustrated in (C). Created with BioRender.com

Here *y_h,asv,s_* is the read count, *β*_0_ is a global intercept, and remaining terms account for technical variation effects in read counts (abundance) as well as biological variation in taxonomic composition. Fixed and random terms dealing with technical variation are as follows: *β*_0_ is the effect of season (*s*) on total read count, u_h_ is a random effect describing variation in total read count among individual hosts (h, where there are multiple samples per host), *u_asv_* is a random effect describing variation in total read count of each ASV across samples and hosts, u_h;s_ is a random effect accounting for library size by describing variation in mean read count in each sample (i.e. host by season), and *u_asv:h:s_* is an additional random effect accounting for row-level variation (over-dispersion). In this example biological effects of interest are specified as follows: *u_asv:h_* is a random effect describing the abundance (read count) of an ASV in host *h, u_asv:s_* is a random effect describing how ASV abundances (read counts) ASV differ between seasons. By apportioning the variance attributed to these different random effects we can assess the relative contributions of these different factors to microbiota composition (Figure 1B-C). For example, a high variance associated with *u_h:s_* would indicate a high degree of technical variation due to library size variation across samples, and high variance associated with *u_asv_* could indicate variation in read counts across ASVs due to over-dispersion introduced by many rare taxa (Figure 1C). With regard to biological inference, variance associated with *u_asv:h_* can be interpreted as indicative of individual ‘repeatability’ of ASV community composition and *u_asv:s_* can be interpreted as reflecting variance associated with compositional shifts across seasons.

Continuing with the above example, we can further use Poisson model outputs to explore which specific taxa are driving differential abundances between groups of interests (e.g. season), which is commonly of great interest in microbiome studies, but for which many existing methods may be affected by library size and normalisation methods (Weiss *et al*. 2015). Differential abundance in this example can be estimated using each ASV-by-season level of the random effect *u_asv:s_* and comparing posterior distributions for each ASV across factor levels. For example, the mean of the posterior distribution for ASV1: summer – that for ASV1: spring can be interpreted as the differential abundance of ASV1 between spring and summer, allowing identification of ASVs which exhibit the largest deviations from the means.

While the Poisson model accounts for variation in library size across samples, there has also been a shift in microbiota research towards explicitly compositional data analysis, which removes any effects of library size (other than in quantifying uncertainty) prior to analysis. The centered log-ratio (CLR) described by Aitchsion (1982) represents one such transformation. We present full details of how to implement the above GLMM approach using CLRs, and then apply this alternative parameterization to the example data described below, in our Supplementary Files.

## 3. A worked example: age and season effects on gut microbiota in wild sheep

To test and illustrate our approach, we obtained faecal samples from Soay Sheep *(Ovis aries)* from the island of Hirta in the St. Kilda archipelago of the Outer Hebrides of Scotland. These animals are free-living and are part of a long-term study in which individuals have been marked and monitored longitudinally since 1985 (Clutton-Brock Pemberton 2004). All animals sampled had been caught and uniquely tagged within a few days of birth, so that their age and sex were known with certainty. Each year, fieldwork teams visit St Kilda in spring to monitor lambing and capture, mark and sample newborn lambs within a few days of birth. Subsequently, each August a larger field team visits to capture, mark and sample animals living in the study area using a series of corral traps (Clutton-Brock & Pemberton 2004).

Two sets of faecal samples were collected, in 2013 and 2016, to allow comparison of the gut microbiota of individuals of different ages (2013) and from the same individuals sampled in different season (2016). The 2013 samples were collected during the August catch and included 30 samples from lambs (around 4 months old) and 28 samples from older adults (ages 2-13). The 2016 samples were collected from a set of 36 females age 1-13 years who were sampled in both spring (around the time of parturition) and then 3-4 months later in August. Microbial DNA was extracted from samples, amplified using bacterial 16S rRNA V4 region primers, and sequenced using the Ilumina MiSeq platform to generate 250 base pair (bp) paired-end reads. Sequences were processed using the DADA2 pipeline in R (v1.12.1) to call amplicon sequence variants (ASVs) (Callahan *et al*. 2016). Full details of sampling, sequencing, and data processing methods are provided in the Electronic Supplementary Material (ESM 1.1-1.4). We conducted standard dissimilarity analysis and PERMANOVA tests on effects of age and season for comparison to the results of our GLMM approach (see ESM 1.4.1 for full details).

### 3.1. Specification of GLMMs

We applied separate GLMMs to the 2013 and 2016 datasets. First, an aggregate dataset for each year was created from the sample metadata, taxonomic classifications for each ASV, and an ASV-by-sample abundance matrix. We fit GLMMS with Poisson errors and log links to each dataset using the Bayesian package ‘MCMCglmm’ (Hadfield 2010) using the approach introduced in Section 2. Models fit to 2013 data, which included samples from hosts of two age classes from a single season (one sample per host) were specified as follows:

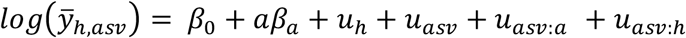

Where *y_h,asv_* is the read count per ASV within each host, *β*_0_ is a global intercept, *β_a_* is the effect of age (*a,* binary factor: lamb versus adult) on total read count, u_h_ is a random effect random effect describing variation in total read count among individual hosts (h, equivalent to sample here where hosts are sampled once each), *u_asv_* is a random effect describing variation in total read count of each ASV across hosts/samples, *u_asv:a_* is a random effect a random effect describing how ASV abundances (read counts) ASV differ between host age classes and *u_asv:h_* is an additional is an additional row-level random effect describing residual variation.

Models fit to 2016 data, which included samples from individual hosts of similar age sampled in both spring and summer of the same year (two samples per host), were specified as follows:

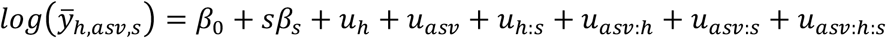

Here *y_h,asv,s_* is the read count, *β*_0_ is a global intercept, *β_s_* is the effect of season (*s*, binary factor: spring versus summer) on total read count, *u_h_* is a random effect describing variation in total read count among individual hosts (h, where there are multiple samples per host), *u_asv_* is a random effect by describing variation in total read count of each ASV across samples and hosts, *u_h:s_* is a random effect describing variation in total read count among individual hosts (h, where there are multiple samples per host), *u_asv:h_* is a random effect describing the abundance (read count) of an ASV in host *h, u_asv:s_* is a random effect describing how ASV abundances (read counts) ASV differ between seasons, and *u_asv:h:s_* is an additional row-level random effect describing residual variation.

*β*_0_ is a global intercept, and remaining terms account for technical variation as well as biological variation of interest. Fixed and random terms dealing with technical variation are as follows: *β*_0_ is the effect of season (*s*) on total read count, u_h_ is a random effect describing variation in total read count among individual hosts (h, where there are multiple samples per host), *u_asv_* is a random effect by describing variation in total read count of each ASV across samples and hosts, u_h;s_ is a random effect accounting for library size by describing variation in mean read count in each sample (i.e. host by season), and *u_asv:h:s_* is an additional random effect accounting for row-level variation (over-dispersion). In this example biological effects of interest are specified as follows: *u_asv:h_* is a random effect describing the abundance (read count) of an ASV in host *h, u_asv:s_* is a random effect describing how ASV abundances (read counts) ASV differ between seasons.

Using this GLMM approach, we calculated the relative contributions to sources of variance in the data from both technical and biological model components. We followed Nakagawa & Schielzeth (Nakagawa & Schielzeth 2013) for calculation of r^2^ from GLMMs with Poisson error distributions. Using this formula, there is a portion of variance equal to 1 minus the sum variance of the model components which represents variance arising from the Poisson distribution. Where multiple samples are present per individual (2016), repeatability of the community composition of ASVs can be estimated as the proportion of variance attributable to differential taxonomic composition across individuals divided by the sum of the variance explained by all other non-technical component terms estimating compositional effects *u_asv:h_*/(*u_asv:s_* + *u_asv:h_* + *u_asv:h:s_*).

We investigated differential abundances as outlined above (2). To extract information of specific bacterial taxa contributing to differential abundance across age groups or season, we used Poisson model outputs and subtracted the posterior distributions for each ASV between group levels (2013: Age; 2016: Season). We used the resultant distribution to calculate a mean difference and highest posterior density interval (HPDI) to estimate differential abundance for each ASV. For example, the mean of the posterior distribution for ASV1:summer – that for ASV1:spring can be interpreted as the differential abundance of ASV between spring and summer, where a difference can be considered significant when credible intervals do not span zero. We calculated p-values of each differential abundance by calculating the proportional overlap of each pairwise set of distributions divided by half the number of stored iterations.

### 3.2. Results

The microbiota communities of Soay Sheep were dominated by two phyla, Firmicutes and Bacteroidetes (Figure S1), as has been previous observed in most vertebrates (Ley *et al*. 2008) Principle coordinates analysis (PCoA) based on Bray-Curtis dissimilarity indicated clustering of samples by age and by season (Figure 2). The result of a PERMANOVA test on the 2013 data set showed a significant difference in group centroids for lambs and adults (pseudo-F = 7.161, p < 0.001), with 11.34% of the variance in gut microbiota composition (R^2^) is explained by differences between lambs and adults. PERMANOVA results for the 2016 data showed that group centroids for April and August are significantly distinct (pseudo-F = 2.026, p = 0.002), but season only explains 2.81% (R^2^) of the observed variance.

**Figure 2.**
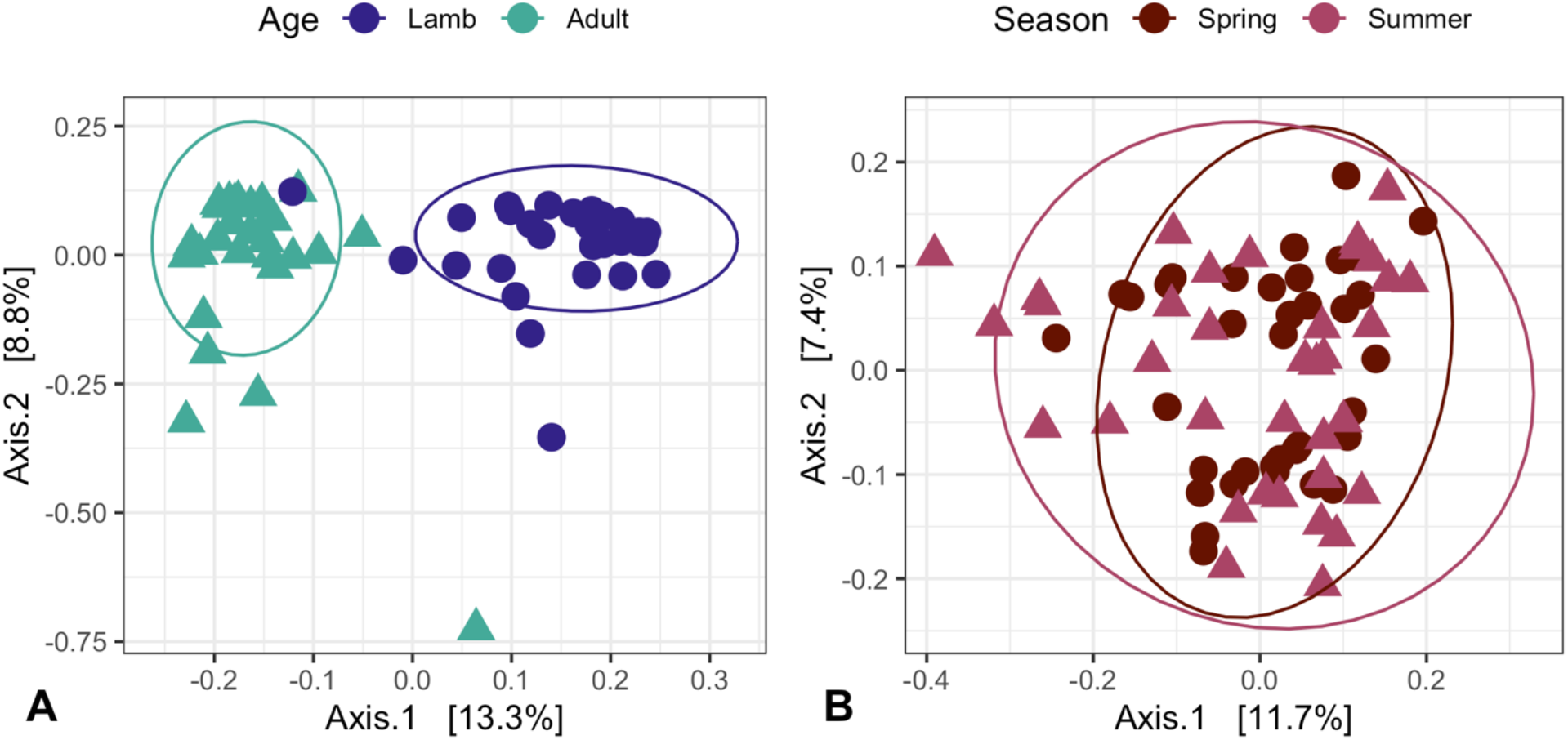
Soay Sheep gut microbiota beta diversity in adults and lambs from 2013 (A) and from April and August of 2016 (B). Principle coordinates (PCoA) plots represent Bray-Curtis dissimilarity indicating clustering of samples by group. Ellipsoids represent a 95% confidence interval surrounding each group.

Poisson GLMMs from 2013 data showed comparable results to ordination approaches, where community composition differed substantially between age classes (Proportion variance *u_asv:a_*: 19.88% CI 18.47-21.39%; Figure 3). Additional effects estimated by the model showed a substantial proportion of variance explained by taxonomic variation in ASV abundance (*u_asv_*: 17.35%), a small portion of variation explained by variation in mean library size across samples (*u_h_* 2013: 1.59 %), and considerable residual variance (estimated by the ‘units’ term in MCMCglmm; *u_asv:h_* 2013: 49.42%; Figure 3).

**Figure 3.**
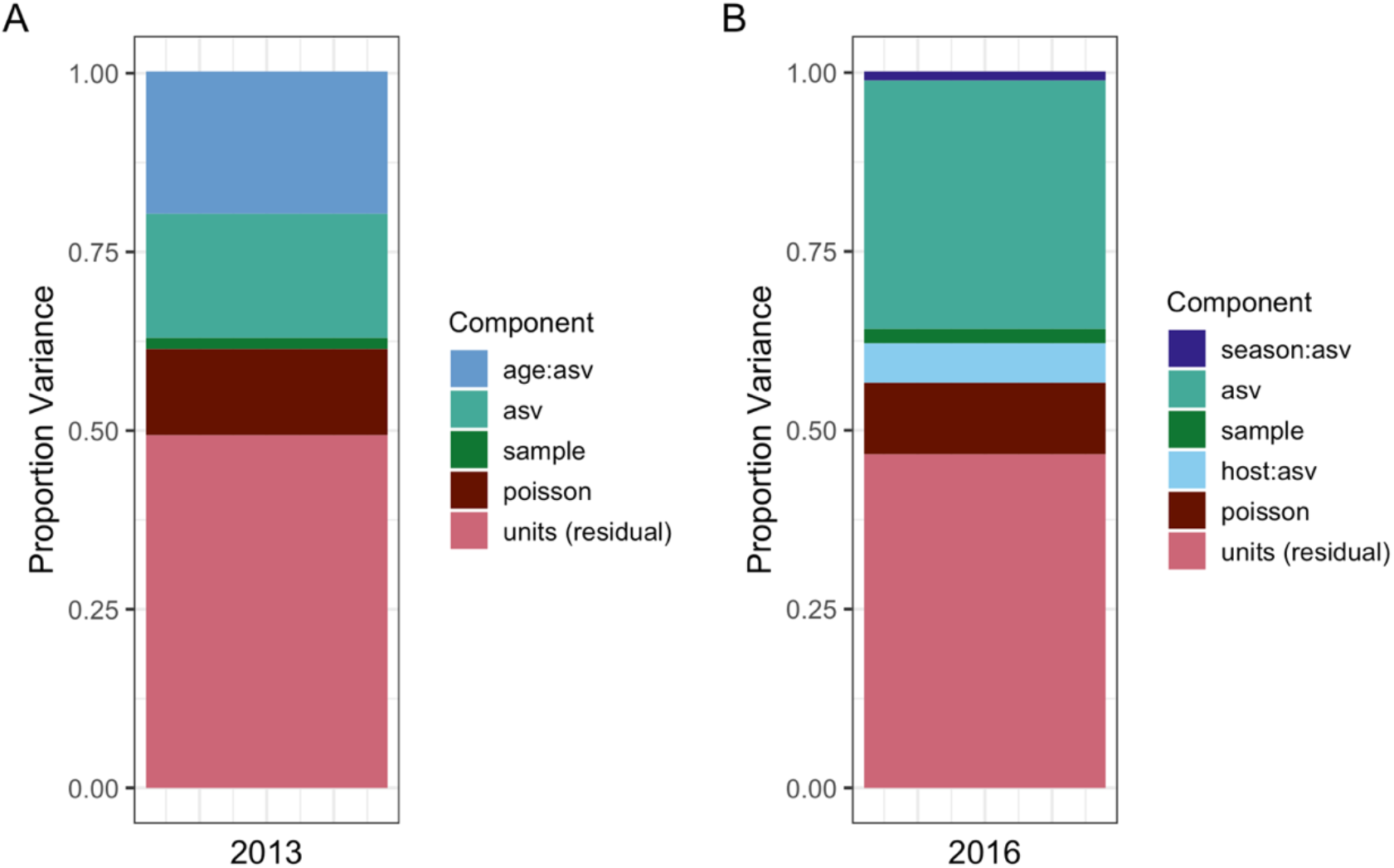
Proportion of variance in bacterial read counts from different ASVs explained by GLMM component terms for two datasets. The 2013 dataset (A) compared gut microbiota across two age classes from individuals sampled once at the same time point), while the 2016 dataset (B) compared samples taken from the same individuals over two seasons.

Poisson GLMMs from 2016 data likewise showed comparable results to ordination approaches, where community composition changed negligibly between seasons (*u_asv:s_*: 1.24% CI 0.99 – 1.44%; Figure 3). The 2016 model showed a substantial proportion of variance explained by taxonomic variation in ASV abundance (*u_asv_* 2016: 34.74%), a small portion of variation explained by variation in mean library size across samples (*u_h:s_* 2013: 1.99 %), and considerable residual variance (*u_asv:h:s_* 2013: 46.67%; Figure 3). Repeated sampling of individuals in 2016 additionally showed moderate variance explained by inter-individual variation in community composition (*u_asv:h_* 2016: 5.49 %). The equated to an individual repeatability of 11.1% for their microbiota community composition across sampling time points.

As outlined in 3.1 we calculated differential abundances using the saved posterior distributions for each random effect level of specific taxa across age classes (2013 data set; Figure 4A & B) and seasons (2016 data set; Figure 4C & D). For the ASV-by-age effect in the 2013 data (*u_asv:a_*, Figure 3), the estimates of taxa-specific differential abundances suggest that ASVs demonstrating significant shifts between lambs and adults belong primarily to two phyla, Firmicutes and Bacteroidetes (Figure 4A-B). 683 out of 2,023 (33.76%) ASVs present in 2013 data showed significant shifts between lambs and adults (50.81% positive shifts, 49.19% negative shifts). Bacteroidetes represented 52.16% of significant positive shifts into adulthood, and Firmicutes represented 72.02% of the significant negative shifts into adulthood (Figure 4B; Table S4). For the ASV-by-season effect in the 2016 data (2016 *u_asv:s_*, Figure 3 & Table S4), very few ASVs (24 of 2,364; 1.02%) showed significant differential abundance between spring and summer sampling (Figure 4C-D; Table S4).

**Figure 4.**
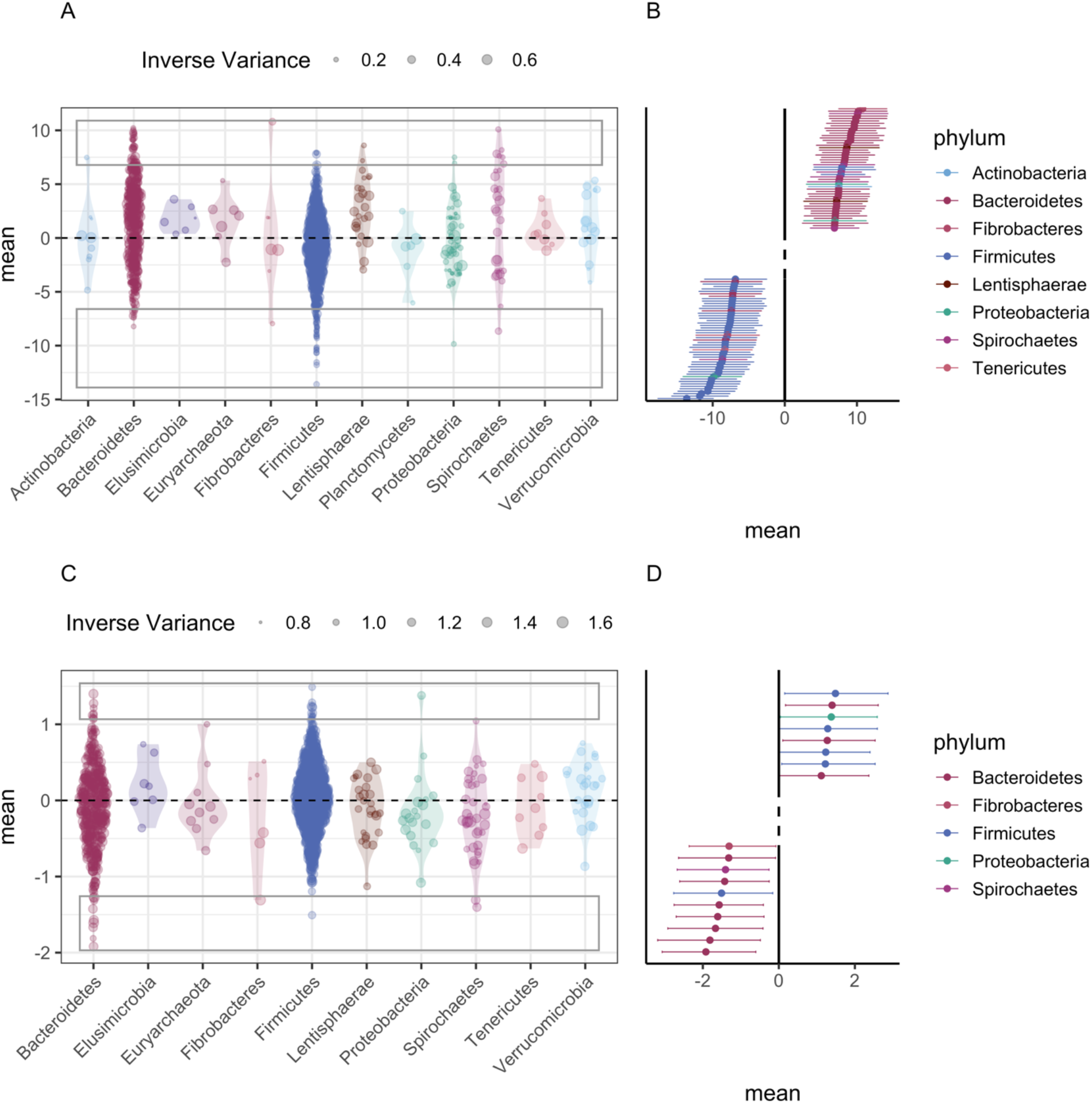
Differential abundances across age classes (A-B) or season (C-D) for individual ASVs calculated from GLMMs with Poisson error families and taxonomic level specified as ASV-only. A & C represent all ASV-level effects. Violin plots represent distribution of effect estimates, size of point represents the inverse variance of the estimate. Rectangles indicate the ASVs with the highest magnitude (positive or negative) differential abundances in Forest plots (B & D). Forest plots represent point estimate and HPDI for the ASVs involved in the 50 (age class) or 20 (season) strongest increases and decreases of abundance.

Our results indicate that there are substantive developmental shifts in the Soay sheep microbiota between lambs and adults, and that the majority of taxa shifting in abundance belong to the Bacteroidetes and Firmicutes. However, this raises the question of whether this shift is because Bacteroidetes are *generally* more abundant in adults and Firmicutes more abundant in lambs, or if the ASVs that show these patterns *just happen* to be in these phyla. To illustrate how GLMMs can be used to address questions of this sort, we modified our models for the 2013 dataset to include additional taxonomic effects of family and phylum, allowing us to identify taxonomic levels most responsible for differential abundances. Details of these phylogenetically more explicit GLMMs and their results and implications are presented in detail in the ESM (see sections 1.4.3, Figure S4, Table S3). Results suggest considerably variation with respect to age across families and ASVs within both the Bacteroidetes and Firmicutes phyla, and that most of the age effects on microbiota community composition occurs at these lower taxonomic levels (Figure S4; Table S3).

## 4. Discussion & wider applications

We have described a novel approach to analysing meta-taxonomic data derived from NGS using a GLMM framework, and have illustrated this method using data describing variation in the gut microbiota community in wild sheep. Our approach represents an important step forward for researchers interested in using meta-taxonomic approaches to understand variation in community structure in complex, non-experimental settings. It allows the well-established power and flexibility of GLMM-based approaches to be harnessed to decompose drivers of variation in NGS-derived data on the taxonomic composition of samples. Our example analyses provide very simple illustrations of how the approach can be used to estimate the contribution of host-related or environmental factors (specifically, age and season) to variation in the community structure of the gut microbiota. We show that results are comparable to those derived from widely-applied ordination-based approaches, and discuss the implications of observed variation in gut microbiota structure with age and season briefly below. However, these analyses are intended mainly as templates to help illustrate the approach, and barely scratch the surface of the types of important outstanding questions the method could be used to tackle with larger-scale data sets. Applying GLMMs to taxonomic-level sequence count data provides a rich toolkit from fields like quantitative genetics to dissect the contributions of different environmental and host factors to variation in community structure, with the potential to advance our understanding of community ecology, host-microbe or -pathogen interactions, and evolutionary dynamics.

In our illustrative analyses of wild Soay sheep, both GLMM and ordination-based approaches identified a stronger effect of age than season on the gut microbiota. Changes in the structure of the gut microbiota across development in early life and during senescence in later adulthood are well-established in human studies (Popkes & Valenzano 2020), but remain poorly understood in natural populations. Our data clearly show the gut microbiota community structure changes between recently weaned lambs and adults, and argues for further longitudinal studies in natural systems to test whether shifts in gut community structure could play a role in patterns of demographic ageing in the wild. A growing number of studies in the wild have documented seasonal gut microbiota changes (Amato *et al*. 2015; Maurice *et al*. 2015; Ren *et al*. 2017; Orkin *et al*. 2019). The absence of a strong seasonal different in our data may be related to the relative homogeneity of the herbivorous diet of Soay sheep, as most previous wild studies are of omnivores with strong seasonal shifts in diet preference. Alternatively, it may be because the spring and summer seasons we sampled in are both periods of relatively high food abundance and quality, and more pronounced change would be observed in autumn and winter when habitat quality and food availability changes more dramatically. In future studies, repeated sampling of the same individuals over time will be crucial to understand the effects of age, environmental and other variables on gut community structure. The application of our approach to such longitudinal metataxonomic data sets will help researchers robustly estimate within-individual patterns of change in community structures over time or space, whilst also estimating how repeatable community structure across hosts.

Our novel GLMM approach allows estimation of key ecological and evolutionary parameters from meta-taxonomic data sets which can advance our understanding of within host-microbe evolutionary dynamics. Individual repeatability of measured phenotypes is an important and widely estimated parameter in ecology and quantitative genetics (Wilson 2018). Estimating the within-host repeatability of microbiota community structure over time can offer insight into the extent to which host control and environmental selection determine species composition (Foster *et al*. 2017). The GLMM structure presented here directly estimates this repeatability across two seasons in our 2016 data set at around 11%, although the small sample size and temporal proximity of the samples should mean we interpret this parameter estimate with caution. However, the model is illustrative and it should be clear that it is readily extendible to mixed-effect model of more a more complex nature to address emerging questions in the field. For example, effects of host relatedness & inbreeding effects on microbiota composition in have been explored previously in microbiome studies but via ordination methods using a small number of genetic clusters as grouping units (Yuan *et al*. 2015). Including host genetic relatedness matrices as a random effect in a mixed-effects model (so-called “animal models”) within our GLMM framework can offer insights into heritability and inbreeding effects on community composition as it compares to other forces in the population (Wilson *et al*. 2010). Factors which can be of both considerable ecological and evolutionary interest, and that can also confound heritability estimates, such as maternal effects or shared nest or litter effects can likewise be incorporated in these models (Kruuk & Hadfield 2007). Statistical advances to address maternal and social effects as well spatial autocorrelation in ecological datasets can also be incorporated in microbiota analyses (Hayward *et al*. 2010; Albery *et al*. 2020). Applied to larger longitudinal data sets, our GLMM approach can allow researchers to directly estimate how different aspects of host state, genotype and environment influence the structure of within-host communities, and address many outstanding questions about the evolutionary and ecological causes and consequences of host-microbe interactions.

Full realisation of the role of microbiota communities in the ecology and evolution of wild organisms depends on both identifying factors with important effects on global microbiota composition and on being able to test whether and how individual taxa or taxonomic groups underpin those effects. Our GLMM approach readily lends itself to addressing both questions. We have illustrated how the approach can be used to identify ASVs involved in community level shifts with age identified in the random effects structure of the 2013 models (Figure 4), and to further decompose the contribution of different taxonomic levels to community level effects (ESM 1.4.3; Table S3). For example, our analysis highlights that analysis at the phylum level could provide a misleading view of compositional shifts associated with age of Soay sheep, and that there is substantial variation at the family level within each phylum (Figure S4). This analysis approach should represent similar insights to linear discriminant analysis (LDA) used in approaches such as LefSE (Segata *et al*. 2011), with the advantage of extraction of this information for multiple factors of interest rather than requirement of a priori knowledge of effects of interest to run differential abundance analysis. Our approach could be further developed beyond the nested taxonomic levels used here to identify taxonomic levels associated with the greatest variance (ESM 1.4.3) to explicitly include the microbiota phylogeny within the GLMMs, allowing environment and host effects on community composition to be estimated accounting more accurately for phylogenetic distances between ASVs (Hadfield & Nakagawa 2010) in a similar manner to UNIFRAC clustering approaches (Lozupone *et al*. 2006). A GLMM-based approach capable of simultaneously dissecting the contributions of host environment, state and genetics alongside microbial phylogeny to variation in microbial community structure seems to us to represent a powerful step towards robustly address emerging questions surrounding the role of the microbiome in ecology and evolution. However, we note however that there are computational challenges associated with incorporating more complex covariance structures into our GLMMs, and provide some discussion of this issue in the supplementary methods with a worked example of an alternate approach (ESM Section 2).

Beyond applications to microbiota community analyses, approaches outlined in this manuscript are applicable more broadly to different types of metataxonomic data being collected across myriad systems and research disciplines. For example, there has been great interest in describing the dynamics of the parasite community as an ecosystem and understanding its influence on host health (Pedersen & Fenton 2007; Graham 2008; Rynkiewicz *et al*. 2015). A growing number of studies apply meta-barcoding to faecal samples to estimate the community structure of gastrointestinal parasite communities (Avramenko *et al*. 2015; Aivelo & Medlar 2017), and our GLMM approach could readily applied to such data sets to dissect the drivers of variation in parasite community composition. Another area of interest within ecology and evolution is using meta-barcoding of faecal samples to estimate diet composition and its relationship to host phenotypes. Bayesian mixed model approaches have also recently been applied to analysis of presence and absence of *Cyanistes caeruleus* (blue tit) diet components and align conceptually with approaches presented in this manuscript (Shutt *et al*. 2020). GLMM approaches to metabarcoding data maintain key similarities to other multivariate community ecology approaches to abundance data (Wang *et al*. 2012) while integrating benefits of ecological and evolutionary approaches to quantifying phenotypic variation. We therefore suggest that approaches presented in this manuscript can be applied across a range of systems and data types for powerful and flexible understanding of complex drivers of community dynamics.

## Acknowledgements

This work was funded by a large NERC grant (NE/R016801/1), and the long-term study on St Kilda was funded principally by responsive mode grants from NERC. We thank Adam Hayward and Jill Pilkington for sample collection, the National Trust for Scotland for support of our work on St Kilda, and QinetiQ and Kilda Cruises for logistical support. We also thank the Ecology Within Team for input in the analysis and manuscript and Josephine Pemberton for support and management of the field project. Figure 1 was created with Biorender.com. Photos on which sheep icons in Figure 1 are based by Hannah Vallin & Martin Stoffel.

## Author contributions

LM and ARS conceived and developed the statistical methods; LM, ARS, AF, HL conducted the analyses; AF, AI, KAW, KW and DHN oversaw and undertook sample collection and laboratory work; ARS wrote the first draft of the manuscript and all authors contributed to writing of the final manuscript.

## Data Accessibility

Data and code for this manuscript will be available on Dryad Repository upon acceptance. Sequences and metadata on which analysis is based can be found on the European Nucleotide Archive (ENA): http://www.ebi.ac.uk/ena/data/view/PRJEB39322

