## Supplementary Material for "A mixed model approach for estimating drivers of microbiota community composition and differential taxonomic abundance"

### Electronic Supplementary Information

$ Joint last authors

### 1. Methods

#### 1.1 Data collection

Faecal samples were collected at at two different time points and each provided a small data set with which to test for differences in the gut microbiota community structure among individuals of different ages and within individuals in different seasons. In August 2013, during the summer catch field season, 58 faecal samples were collected rectally from animals at capture. 30 of these samples were taken from lambs (N = 15 Male, 15 Female) and 28 were taken from adults (age range 2 – 13 years; N = 6 Male, 22 Female). In 2016 faecal samples were collected from pasture from 36 marked adult female sheep (age range 1- 13 years) in spring (between 4th and 27th April). These samples were collected by observing identifiable, marked individuals at close quarters until they defecated and then collecting the faecal samples from the pasture within a minute. The same individuals were resampled during the summer catch around three months later (12th – 18th August) using rectal faecal collection. Samples were stored at -20C within approximately 3 hours of collection.

#### 1.2 Laboratory processing

DNA extractions for the 2013 samples were carried out using the Qiagen 'QIAamp DNA Stool' kit. DNA extractions for the 2016 samples were carried out using the MoBio PowerFecal DNA Isolation kit (Mo Bio, Carlsbad, CA, USA) following the manufacturers guidelines. DNA extractions were stored at -20C until they were amplified using polymerase chain reaction. The V4 region of the bacterial 16S rRNA gene was amplified using the 515F forward primer and Golay barcoded 806R reverse primer series (Caporaso et al, 2012). Next-generation DNA sequencing was carried out at Source BioScience LifeSciences for the 2013 samples and at Edinburgh Genomics for the 2016 samples using Illumina MiSeq v2 to generate 250 base pair (bp) paired-end reads.

##### 1.2.1 2013 Samples

Adaptation to the standard Qiagen Stool DNA extraction protocol: All centrifugation steps were carried out at 15,000 rcf/g, Step 4 – Centrifuge for 1.5 minutes, Step 5 – Supernatant volumes below 1.2ml were supplemented with Buffer ASL, Step 7 & 8 – Centrifuge for 4.5 minutes, Step 12 – Incubate at 95C for 10 minutes, Step 14 & 15 – Centrifuge for 1.5 minutes, Step 16 – Centrifuge for 4.5 minutes. All PCRs were performed in 25ul reactions: 12.25ul $H_2O$, 5ul Q Solution, 2.5ul PCR Buffer (10x), 1ul dNTP Mix, 0.25ul Hot star taq polymerase, 1ul Forward Primer (10uM), 1ul Reverse Primer (10uM) and 2ul DNA. The PCR conditions were as follows: Initial denaturation at 96C for 15 minutes then 35 cycles of denaturing at 94C for 45 seconds, annealing at 50 degrees for 60 seconds, extension at 72C for 90 seconds and a final extension at 72C for 10 minutes. PCR products were cleaned using Zymo DNA 96 Clean and Concentrator kit (all carried out by Source BioScience LifeSciences).

##### 1.2.2 2016 Samples

All PCRs were performed in 25ul reactions using Roche reagents: 18.5ul $H_2O$, 1ul $MgCl_2$(25mM), 2.5ul PCR Buffer (10x), 0.5ul dNTP Mix (10mM), 0.25ul Taq DNA polymerase (5U/ul), 0.625ul Forward Primer (10uM), 0.625ul Reverse Primer (10uM) and 1ul DNA. The PCR conditions were as follows: Initial denaturation at 94C for 3 minutes then 25 cycles of denaturing at 94C for 45 seconds, annealing at 50 degrees for 60 seconds, extension at 72C for 90 seconds and a final extension at 72C for 10 minutes. PCR products were cleaned up using Wizard SV Gel and PCR Clean-up System (reference).

#### 1.3 Data Processing: DADA2 Pipeline Parameters

Sequences were processed using the DADA2 pipeline in R (v1.12.1) to call amplicon sequence variants (ASVs) (Callahan *et al.* 2016). Briefly, forward and reverse reads were examined for quality, trimmed and filtered (See Supplementary Material). Sequences for both datasets were independently denoised using the DADA2 algorithm (Callahan et al 2016), forward and reverse reads merged, ASVs inferred and tabulated, and putative chimeras removed. Taxonomy was assigned to each ASV using the naive Bayesian classifier of DADA2 and the Silva Project v132 database formatted for DADA2.

Bioinformatics processing followed the workflow of the DADA2 Pipeline Tutorial (1.12) (available at https://benjjneb.github.io/dada2/tutorial.html). Sequences for both datasets were examined for quality to determine trimming parameters. Sequences were trimmed at 240bp (Forward) and 200bp (Reverse) in 2016, and at 200bp (Forward) and 150bp (Reverse) in 2013, where sequences were of poorer quality. Standard filtering parameters were used (maxN=0, truncQ=2, rm.phix=TRUE and maxEE=2), with the exception of an increase of maxEE=10 for 2013 sequences to improve read retention during filtering. The average proportion of reads retained per sample at the end of the bioinformatics processing was ~0.7 for 2016 data and ~0.6 for 2013 data.

Sample metadata, taxonomy tables, and an ASV abundance matrix were integrated into a phyloseq object for downstream analysis (McMurdie & Holmes 2013). 2013 data contained 5,156 ASVs and 2,454,172 total reads (average 42,313.31; range 23,462-56,289). 2016 phyloseq objects contained 8,593 ASVs and 3,911,335 total reads (average 42,313.31; range 32,940-103,743). Prior to analysis an abundance filter was applied to datasets of both years, eliminating ASVs with a total abundance of <100 reads, resulting in 2,023 ASVs for 2013 and 2,364 for 2016. We ran a series of mixed models using multiple filtering criteria to ensure model outputs were not dependent on filtering threshold sensitivity (see Supplementary Material).

#### 1.4 Statistical analysis

##### 1.4.1 Traditional analysis additional tests

All analyses were conducted in R version 3.6.1 (R Core Team 2019). To provide a comparison with more traditional statistical approaches to meta-barcoding data, we normalised read abundances to compositional proportion data and calculated pairwise dissimilarities among samples using Bray-Curtis in the Phyloseq package of R (McMurdie & Holmes 2013). To assess the extent to which age and season predicted microbiota composition these dissimilarities were used in principle coordinates analysis (PCoA). Permutational analysis of variance (PERMANOVA) was used to study the difference in group means and was carried out using the Vegan function 'adonis' (Anderson 2001).
In addition to beta diversity, alpha diversity (diversity within an individual) was calculated using the Simpson index. This was calculated across categorical variables: age and season and homogeneity of dispersion was tested using a PERMDISP with the function 'betadisper' in the Vegan package (Anderson, 2001; Oksanen et al, 2013).

##### 1.4.2 Poisson GLMM Abundance Threshold Validation

As a validation of data filtering in our mixed model approach, we ran 2016 models on data using the following criteria: (i) no abundance filter, (ii) a threshold of 50 total reads, (iii) a threshold of 100 total reads, and (iv) a threshold of 500 total roads to ensure model outputs were not dependent on data processing steps. Neither the rank order nor the absolute values of proportion variance explained by component terms of interest were notably affected by abundance filtering levels (Figure S2). The primary effect of increased abundance filtering was a reduction in variance explained by distributional Poisson variation in the data, where and the highest variance explained by the row-level residual variance of differential abundance across samples (component "units", Figure S2).

##### 1.4.3 Taxonomic resolution models

We fit a set of models to further investigate the drivers of differential abundance across host (age) and environment (season) groups by including multiple levels of taxonomy (ASV, family, phylum). These models were fit as GLMMs with Poisson error families for both 2013 and 2016 as described above, with additional random effects describing family and phylum effects as follows for 2013. Resultant model structure each dataset were as follows:

2013:

$$log(\bar{y}_{h,asv}) = \beta_0 + a\beta_a + u_h + u_{asv} + \boldsymbol{u_f} + \boldsymbol{u_p} + u_{asv:a} + \boldsymbol{u_{f:a}} + \boldsymbol{u_{p:a}} + u_{asv:h}$$

2016:

$$log(\bar{y}_{h,asv,s}) = \beta_0 + s\beta_s + u_{h:s} + u_{asv} + \boldsymbol{u_f} + \boldsymbol{u_p} + u_{asv:s} + u_{asv:s} + \boldsymbol{u_{f:s}} + \boldsymbol{u_{p:s}} + u_{asv:h:s}$$

Where bolded terms represent additional terms incorporated to models presented in the main text (Section 3.1). In each model, $\boldsymbol{u_f}$ and $\boldsymbol{u_p}$ represent taxonomic variation in overall abundance at the family and phyla levels, and family- or phyla- level variation in community composition structure across age classes ($\boldsymbol{u_{f:a}} + \boldsymbol{u_{p:a}}$) or seasons ($\boldsymbol{u_{f:s}} + \boldsymbol{u_{p:s}}$) are described by additional interaction terms. Differential abundance calculations using posterior distributions were carried out for the family and phylum level as described in the main text for differential abundance of ASVs.

This modified model for 2013 data showed that that among the 20% overall variance associated with compositional differences between age classes (Figure 3), 13% is attributable to variance at the sequence variant (ASV) level, 5% at the family level, and 2% at the phylum level (Table S3). Low proportion of variance at the phylum level when multiple taxonomic levels are taken into account is due to variation in the direction of ontogenic shifts at the family and ASV levels even within the two phyla most associated with differential ASV abundance (Figure 5). For example, although two families within the Bacteroidetes do show positive shifts into adulthood, some show no difference across age classes, and one (Bacteroidaceae) shows a negative shift (Figure 5). Similarly, although Firmicutes primarily are implicated in negative significant shifts in GLMMs accounting only for ASV (Figure 4A-B), models decomposing the family and phylum levels show that only one family is associated with a significant shift (Figure 5). These results illustrate how GLMMs can be used to decompose the taxonomic levels at which ecological effects occur in microbiota communities

#### 2. A Centred log-ratio (CLR) approach

##### 2.1 Background

The CLR takes the form of the log-ratio of the count for a given taxon in a given sample and the geometric mean of the counts of all taxa in that sample. The CLR can be calculated for any taxon *i* in a sample of *n* taxa as:

$$clr_i = log\left(\frac{x_i}{G(x)}\right)$$

$$G(x) = \sqrt[n]{x_1 \times x_2 \times \dots x_n}$$

The CLR has several desirable properties. Firstly, as a log-ratio it is approximately Gaussian in distribution, allowing the use of a Gaussian linear mixed model. Secondly, the mean of the CLR of all taxa in any sample will be zero, allowing for simplification of our random effects structure. Using the CLR we can for example explore the effect of season on microbiota composition using the following Gaussian LMM:

$$clr_{h,asv,s} = \beta_0 + u_{asv} + u_{asv:h} + u_{asv:s} + u_{asv:h:s}$$

Note that here the random effects not including ASV as an element are removed as the mean of the CLR within samples, and therefore across hosts and seasons, is zero. This reveals the role of the different random effects here – the effects $u_{asv}$, $u_{asv:h}$, $u_{asv:s}$ and $u_{asv:h:s}$ describe effects on microbiota composition, while the effects $u_h$ and $u_{s,h}$ are effects on the mean read count across all taxa. Finally, the three terms $u_{asv:h}$, $u_{asv:s}$ and $u_{asv:h:s}$ and their sizes relative to each other are of most biological interest. These three terms capture how much the abundance of a taxon are affected by consistent host effects, seasonal effects, and idiosyncratic effects (the row level effect of seasonal effects varying by host). These effects capture changes in a taxon's abundance across hosts and seasons, after controlling for the mean abundance of each taxon, sample, and season.

Although we used GLMMs with Poisson error distributions to illustrate the benefits of a GLMM approach, we also acknowledge several limitations with this method. First, even applied to illustrative datasets with a modest number of random effects, run-time for MCMCglmm-based Poisson models is very slow due to the high number of rows for reads-by-ASV-by-sample data frames. We therefore anticipate that dependent on sample size, diversity of ASVs present, and complexity of covariates required these models may become prohibitively slow. For this reason, we explored gaussian models using CLR as responses here as an alternative approach. CLR transformations are increasingly common, acknowledge the compositional nature of count data, allow estimation of error associated with CLR estimates for each ASV per sample, and allow gaussian error distributions to be fit in models structurally similar to Poisson models (Gloor *et al.* 2017).

#### 2.2 Centered log-ratio approach applied to sheep data

We used the package 'ALDEx2' (Fernandes *et al.* 2013) to calculate a centered log-ratio for each asv in each sample. In addition to clr values, ALDEx2 estimates technical variation for each asv from a probability distribution using Monte-Carlo instances drawn from the Drichlet distribution, which maintains the compositional nature of the data. MCMCglmm allows specification of this technical variance as measurement error within model syntax. The variance for each asv:sample observation was therefore calculated from probability distributions generated from 128 Monte-Carlo instances and was specified as the measurement error for the response variable in CLR models. CLR model syntax to estimate the same terms as Poisson GLMMs (Main text Section 3.1), with the exclusion of the effects estimating variation in mean library size across age $a\beta_a$ (2013) or season $s\beta_s$ (2016) and across sample $u_h$ (2013) or $u_{h:s}$ (2016), given that this technical variation is contained within the measurement error structure described above. Model syntax for ratio-normalised values were therefore as follows:
2013:

$$clr_{h,asv} = \beta_0 + u_{asv} + u_{asv:a} + u_{asv:h}$$

2016:

$$clr_{h,asv,s} = \beta_0 + u_{asv} + u_{asv:h} + u_{asv:s} + u_{asv:h:s}$$

Models were specified in MCMCglmm with gaussian error families and proportion of variance explained by each model component was calculated as described in main text (Section 3.1)

#### 2.3 Centered log-ratio results

GLMMs using CLR normalised values as the response variable showed agreement with both traditional approaches and raw read count models (Figure S3). Proportion of variance attributable to differential community composition across age classes ($u_{asv:a}$, 11.0%) was substantial but more conservative compared to Poisson models, while proportional variance for season effects ($u_{asv:t}$, 0.8%) was comparable to Poisson models. As in Poisson models, taxonomic variation in ASV abundance explained a high proportion of variance ($u_{asv}$; 2013: 28.0%, 2016: 44.0%); differential taxonomic abundances across individuals ($u_{asv:h}$) explained 4.0% amount of variance in 2016 (Figure S3). Finally, technical variation estimated as measurement error in CLR models explained a moderate proportion of variance in both years (mev; 2013: 9.9%, 2016: 8.3%).

Overall, CLR models applied to illustrative datasets here showed comparable results to Poisson models with regard to proportion variance attributable to model components. There was slight divergence in the relative proportions of variance explained by CLR vs Poisson models in that the age effect was associated with a slightly more modest proportion variance in CLR models, which may be attributable to the transformations accounting for the limitations of the count data as bound by upper limits of the sequencing platform. We suggest that CLR transformations therefore can offer a more computationally-efficient route for GLMM approaches without sacrificing model complexity. This may be particularly useful for instances where it is desirable to extract posterior distributions to estimate differential abundance of specific taxa given storage of posterior distributions for all random effect levels introduces significant demands on RAM. This possible issues may be circumvented by exploring packages beyond MCMCglmm including ASREML (Gilmour *et al.* 2009) for gaussian models which estimates mean and 95% confidence intervals (rather than saving posterior distributions) for random effects or brms (Bürkner 2017) which uses Hamiltonian sampling rather than Markov chain Monte Carlo methods and is more amenable to parallelisation. We believe there are several relevant options given the significant interest for understanding functional implications of host microbiota community composition.

#### 3. Supplementary Figures & Tables

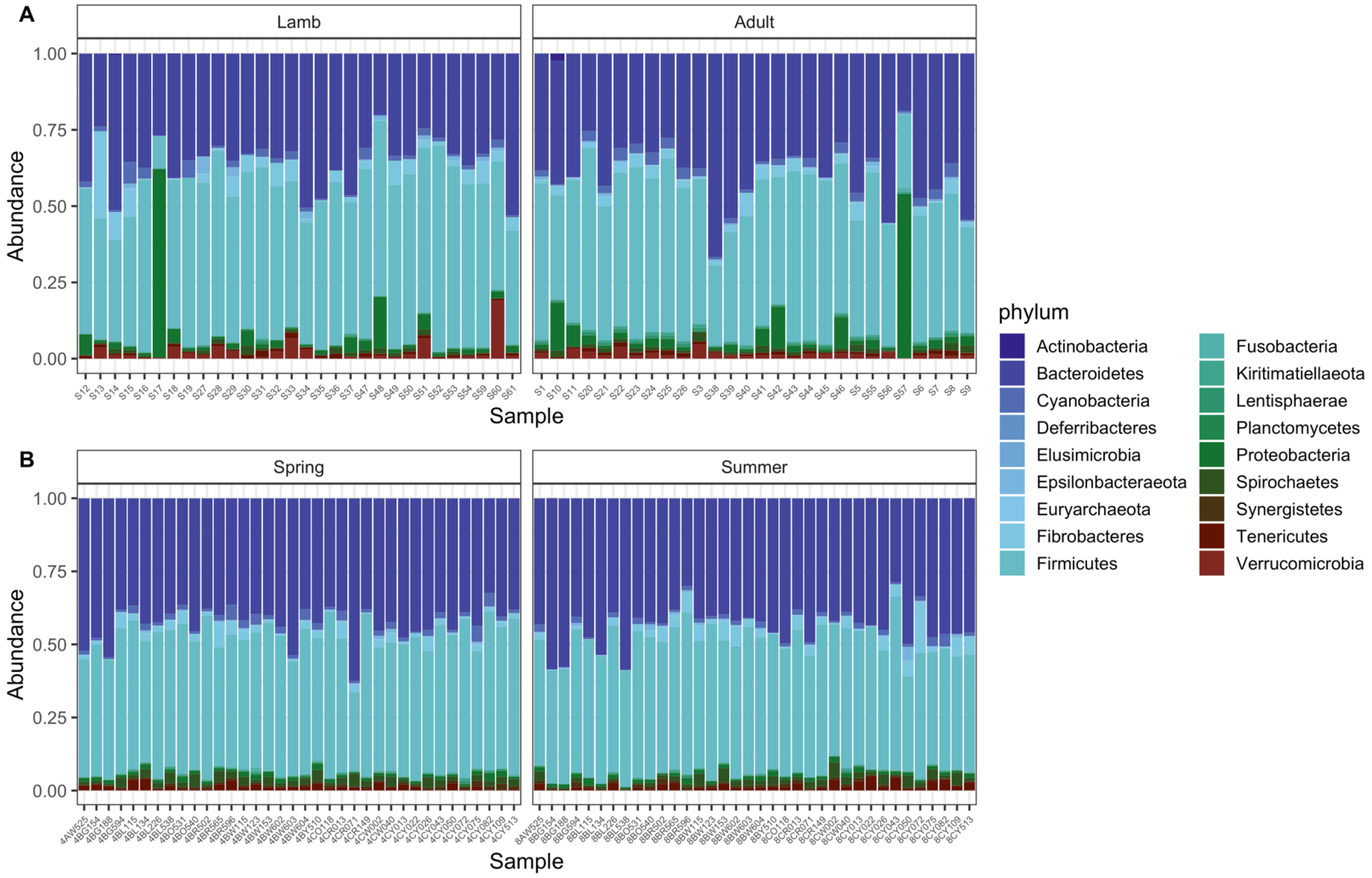


**Figure S1**. Relative abundance of bacterial phyla present in Soay Sheep data from (A): 2013 and (B): 2016.

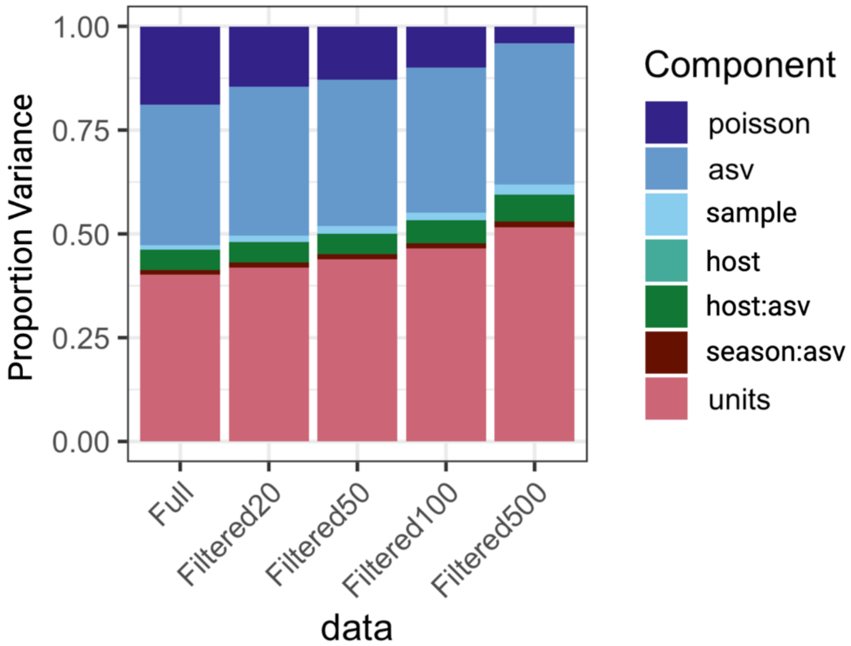


**Figure S2.** Proportion variance of GLMM component terms across multiple levels of initial abundance filtering.

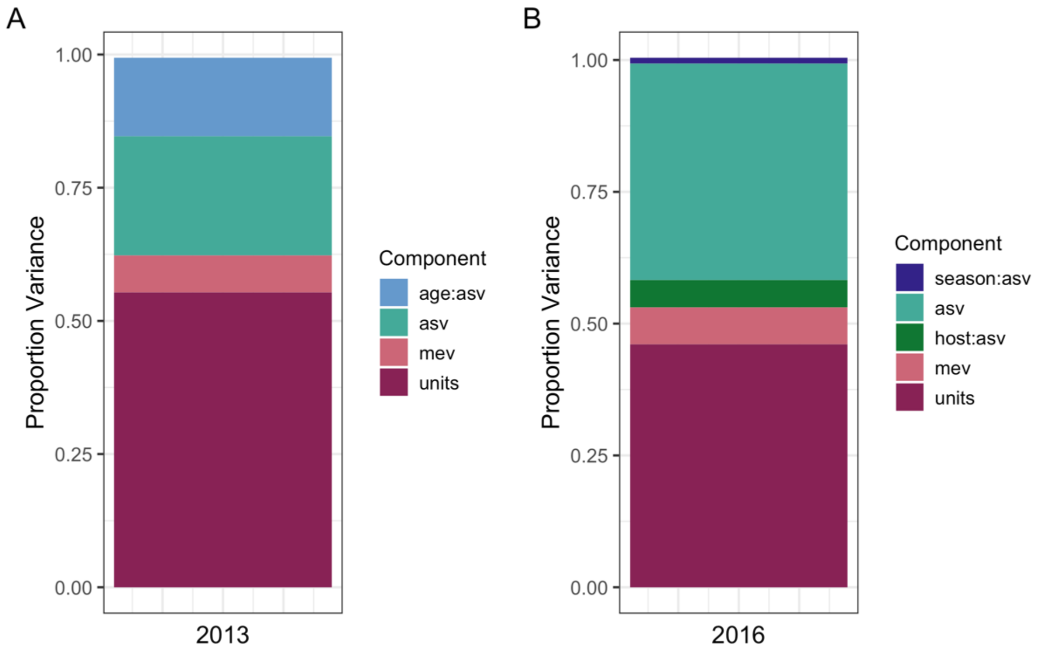


**Figure S3.** Proportion variance of GLMM component terms for gaussian models using ratio-transformations for (A) 2013 data and (B) 2016 data.

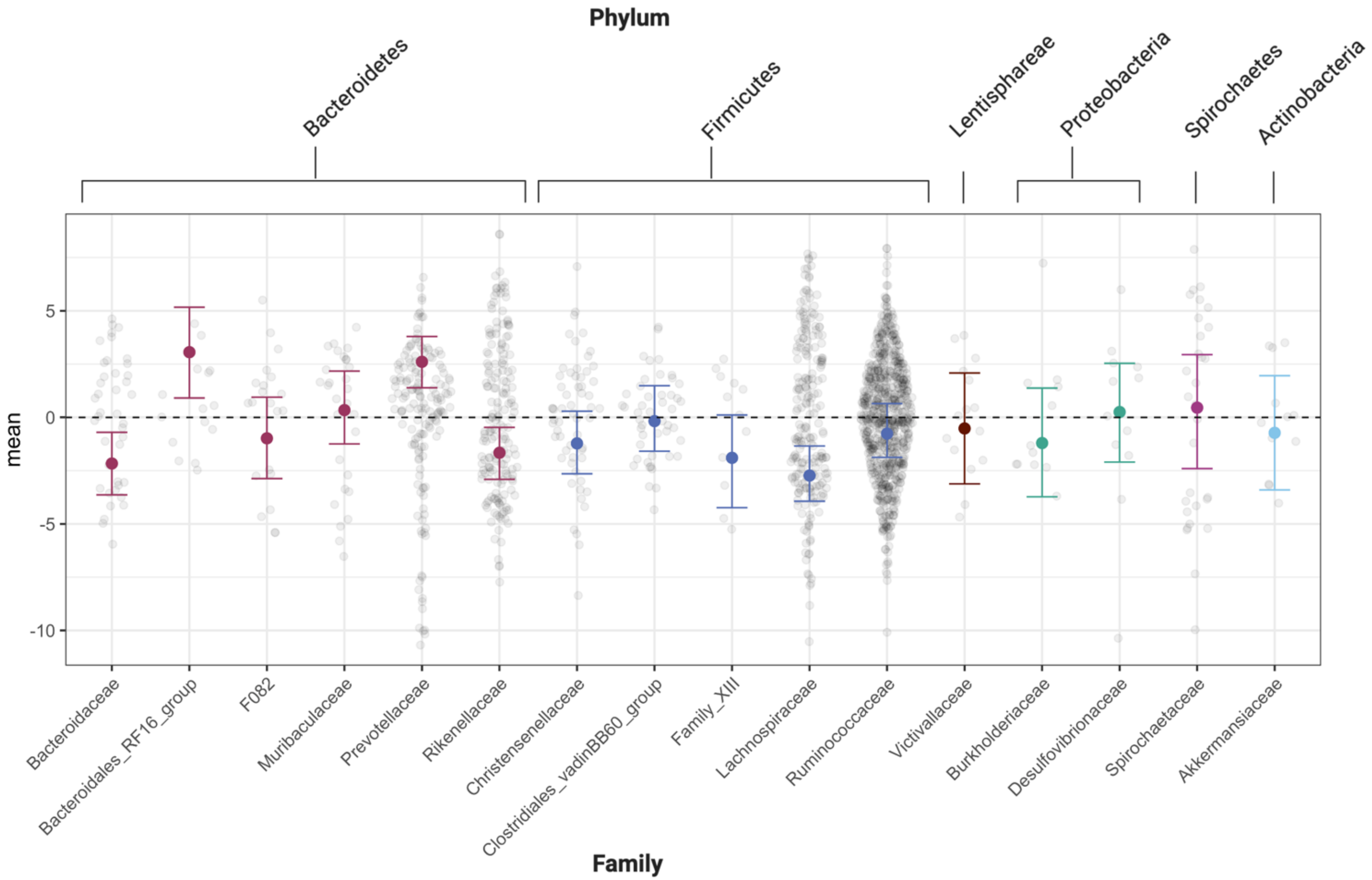


**Figure S4.** Differential abundances across age classes accounting for multiple levels of taxonomy. Data represents 2013 model estimates represent from GLMMs with Poisson error families and taxonomic level specified as ASV, family, and phylum. Points in sina distributions represent ASV-level estimates, point and errorbars represent mean and HPDI for family-level estimates, and colours indicate phylum categories.

**Table S1**. MCMCglmm output for Poisson GLMMs. Blank cells indicate NAs.

| dataset | variable | post.mean | conf.low | conf.high | eff.samp | pMCMC | effect | propVariance | lHPD | uHPD |
|---|---|---|---|---|---|---|---|---|---|---|
| 2013 | Intercept | -5.458 | -5.819 | -5.120 | 1000.000 | 0.001 | fixed | | | |
| | ageAdult | 1.126 | 0.652 | 1.637 | 839.332 | 0.001 | fixed | | | |
| | sample | 0.725 | 0.443 | 1.014 | 1000.000 | | random | 0.0159 | 0.0110 | 0.0244 |
| | asv | 7.336 | 6.479 | 8.088 | 1000.000 | | random | 0.1735 | 0.1552 | 0.1885 |
| | age:asv | 8.551 | 7.807 | 9.266 | 860.142 | | random | 0.1988 | 0.1847 | 0.2139 |
| | units (id:asv) | 21.132 | 20.729 | 21.619 | 981.480 | | residual | 0.4941 | 0.4845 | 0.5086 |
| 2016 | Intercept | -2.651 | -2.918 | -2.359 | 1000.000 | 0.001 | fixed | | | |
| | seasonSummer | 0.066 | -0.258 | 0.392 | 1048.847 | 0.684 | fixed | | | |
| | sample | 0.588 | 0.408 | 0.807 | 1000.000 | | random | 0.0200 | 0.0146 | 0.0287 |
| | asv | 9.547 | 8.969 | 10.142 | 1000.000 | | random | 0.3474 | 0.3317 | 0.3611 |
| | id:asv | 1.491 | 1.285 | 1.670 | 1027.140 | | random | 0.0549 | 0.0468 | 0.0611 |
| | season:asv | 0.332 | 0.274 | 0.396 | 1000.000 | | random | 0.0124 | 0.0099 | 0.0144 |
| | Units (id:asv:season) | 12.792 | 12.530 | 13.054 | 1000.000 | | residual | 0.4668 | 0.4525 | 0.4779 |

**Table S2**. MCMCglmm output for CLR Gaussian GLMMs. Blank cells indicate NAs.

| dataset | variable | post.mean | conf.low | conf.high | eff.samp | pMCMC | effect | propVariance | lHPD | uHPD |
|---|---|---|---|---|---|---|---|---|---|---|
| 2013 | Intercept | 0.889 | 0.802 | 0.984 | 1000.000 | 0.001 | fixed | | | |
| | age:asv | 2.136 | 1.977 | 2.281 | 942.252 | | random | 0.147 | 0.139 | 0.160 |
| | asv | 3.286 | 2.984 | 3.575 | 782.964 | | random | 0.224 | 0.213 | 0.246 |
| | mev | 1.000 | 1.000 | 1.000 | 0.000 | | random | 0.069 | 0.068 | 0.071 |
| | units (id:asv) | 7.935 | 7.848 | 8.031 | 1000.000 | | residual | 0.554 | 0.540 | 0.564 |
| 2016 | Intercept | 1.025 | 0.916 | 1.113 | 1000.000 | 0.001 | fixed | | | |
| | asv:id | 0.731 | 0.630 | 0.809 | 1000.000 | | random | 0.0517 | 0.0444 | 0.0569 |
| | season:asv | 0.150 | 0.122 | 0.177 | 809.734 | | random | 0.0108 | 0.0084 | 0.0124 |
| | asv | 5.786 | 5.457 | 6.112 | 1000.000 | | random | 0.4105 | 0.3939 | 0.4220 |
| | mev | 1.000 | 1.000 | 1.000 | 0.000 | | random | 0.0701 | 0.0688 | 0.0721 |
| | units (id:asv:season) | 6.531 | 6.429 | 6.641 | 1000.000 | | residual | 0.4609 | 0.4475 | 0.4733 |

**Table S3**. MCMCglmm output for Poisson GLMM with nested taxonomic structure for 2013 age effects. Blank cells indicate NAs.

| dataset | variable | post.mean | conf.low | conf.high | eff.samp | pMCMC | effect | propVariance | lHPD | uHPD |
|---|---|---|---|---|---|---|---|---|---|---|
| 2013 | Intercept | -6.659 | -7.645 | -5.744 | 1000.000 | 0.001 | fixed | | | |
| | ageAdult | 2.590 | 1.307 | 3.637 | 1000.000 | 0.001 | fixed | | | |
| | sample | 0.960 | 0.549 | 1.445 | 1000.000 | | random | 0.017 | 0.012 | 0.031 |
| | asv | 7.748 | 6.839 | 8.486 | 1000.000 | | random | 0.158 | 0.144 | 0.178 |
| | age:asv | 6.427 | 5.786 | 7.038 | 1000.000 | | random | 0.133 | 0.121 | 0.148 |
| | phylum | 0.289 | 0.000 | 1.099 | 844.192 | | random | 0.000 | 0.000 | 0.023 |
| | sample:phylum | 0.967 | 0.692 | 1.217 | 1000.000 | | random | 0.018 | 0.015 | 0.025 |
| | age:phylum | 1.190 | 0.168 | 2.504 | 970.971 | | random | 0.019 | 0.004 | 0.051 |
| | family | 0.587 | 0.000 | 1.920 | 608.939 | | random | 0.000 | 0.000 | 0.040 |
| | sample:family | 2.221 | 1.892 | 2.570 | 1000.000 | | random | 0.048 | 0.040 | 0.055 |
| | age:family | 2.597 | 1.185 | 4.243 | 733.308 | | random | 0.054 | 0.024 | 0.084 |
| | units (id:asv) | 19.419 | 19.014 | 19.797 | 863.139 | | residual | 0.408 | 0.384 | 0.429 |

**Table S4.** Representation across phyla of significant ASV-level compositional shifts. Blank cells indicate that the phyla had no ASVs with significant abundance shifts.

| | Age: Adults v Lambs | | | | Season: Summer v Spring | | | |
|---|---|---|---|---|---|---|---|---|
| | Positive Shifts | | Negative shifts | | Positive Shifts | | Negative shifts | |
| phylum | n ASV | % total ASVs | n ASV | % total ASVs | n ASV | % total ASVs | n ASV | % total ASVs |
| Bacteroidetes | 181 | 52.16 | 62 | 18.45 | 3 | 37.50 | 9 | 56.25 |
| Firmicutes | 89 | 25.65 | 242 | 72.02 | 4 | 50.00 | 1 | 6.25 |
| Kiritimatiellaeota | 14 | 4.03 | 1 | 0.30 | | | 1 | 6.25 |
| Cyanobacteria | 13 | 3.75 | 6 | 1.79 | | | 2 | 12.50 |
| Spirochaetes | 13 | 3.75 | 7 | 2.08 | | | 2 | 12.50 |
| Lentisphaerae | 12 | 3.46 | | | | | | |
| Proteobacteria | 9 | 2.59 | 3 | 0.89 | 1 | 12.50 | | |
| Tenericutes | 6 | 1.73 | 12 | 3.57 | | | | |
| Verrucomicrobia | 5 | 1.44 | | | | | | |
| Actinobacteria | 1 | 0.29 | 1 | 0.30 | | | | |
| Elusimicrobia | 1 | 0.29 | | | | | | |
| Euryarchaeota | 1 | 0.29 | | | | | | |
| Fibrobacteres | 1 | 0.29 | 1 | 0.30 | | | 1 | 6.25 |
| Synergistetes | 1 | 0.29 | | | | | | |
| Planctomycetes | | | 1 | 0.30 | | | | |

#### Supplementary Material References


Anderson, M.J. (2001). A new method for non-parametric multivariate analysis of variance. *Austral Ecology*, 32–46.

Bürkner, P.-C. (2017). brms: An R Package for Bayesian Multilevel Models Using Stan. 1–1.

Callahan, B.J., McMurdie, P.J., Rosen, M.J., Han, A.W., Johnson, A.J.A. & Holmes, S.P. (2016). DADA2: High-resolution sample inference from Illumina amplicon data. *Nature methods*, **13**, 581–583.

Fernandes, A.D., Macklaim, J.M., Linn, T.G., Reid, G. & Gloor, G.B. (2013). ANOVA-Like Differential Expression (ALDEx) Analysis for Mixed Population RNA-Seq (J. Parkinson, Ed.). *PLoS One*, **8**, e67019.

Gilmour, A.R., Gogel, B.J., Cullis, B.R. & Thompson, R. (2009). ASReml User Guide. 1–398.

Gloor, G.B., Macklaim, J.M., Pawlowsky-Glahn, V. & Egozcue, J.J. (2017). Microbiome Datasets Are Compositional: And This Is Not Optional. *Frontiers in microbiology*, **8**, 2224.

McMurdie, P.J. & Holmes, S. (2013). phyloseq: An R Package for Reproducible Interactive Analysis and Graphics of Microbiome Census Data (M. Watson, Ed.). *PLoS One*, **8**, e61217.
